# Regional Similarity to Sex-Typical Young Adult Brain Architecture is a Longitudinal Correlate of Neuroinflammation and Depression in Later Adulthood

**DOI:** 10.64898/2026.10.04.756560

**Authors:** Raluca Petrican, Benedict D. Michael, Erin Mulderig

## Abstract

Lifespan trajectories of psychiatric and neurodegenerative risk show robust overlap and marked sex differences. Nonetheless, their neurobiological underpinnings are still poorly understood. One potential culprit is the predominant categorical definition of sex, which overlooks the tapestry of sex-dependent and sex-independent features recently shown to typify each human brain. Shifting from a holistic, dichotomous approach to regionally specific, continuous assessments of sex differences in brain function could help identify individualized therapeutic targets for both psychiatric and neurological conditions. To probe the psychiatric and neurological relevance of functional alignment with sex-typical brain architecture, we analyzed longitudinal neuroimaging and clinical data from a healthy aging cohort (N = 353, aged 36-87 years). We identified complementary patterns of deviance from sex-typical brain architecture along the canonical sensorimotor-to-association axis, which overlapped areas enriched in microglial genes and genes regulating sex steroid metabolism. The two connectivity patterns differentiated longitudinally between psychiatric symptoms and neurodegeneration as a function sex and age: one connectivity pattern related to more severe future depression in younger women, whereas the second related to later neurodegeneration (higher GFAP and pT217 blood concentration) and poorer fluid cognition (inhibition, episodic memory, processing speed, working memory) in older men. The aforementioned effects showed some context-specificity, as they involved connectivity patterns characterized during awake rest and cognitive control, but not associative memory performance. Taken together, our findings attest to the value of integrating regionally specific, continuous assessments of sex differences in brain function with inflammation-relevant biomarkers in order to support precision medicine approaches across the life course.

---

Sex-dependent trajectories of lifespan neurobiological development have been implicated in the emergence of sex-differential profiles of psychiatric and neurodegenerative risk (Bangasser and Cuarenta, 2021; Dhamala et al., 2023; Eme, 2016; Gilmore et al., 2018). Mounting evidence points to the foundational role of gonadal hormones in the maturation of brain regions that are sexually dimorphic across species and underpin the female-biased risk for internalizing disorders and the male-biased vulnerability to externalizing disorders (Ahmed et al., 2008; Pfeifer and Allen, 2021; Piekarski et al., 2023; Solmi et al., 2022). Gonadal hormones have also been implicated in both normative and pathological brain aging (Barth et al., 2015; Lopez-Lee et al., 2024; Newhouse and Dumas, 2015), with more severe neurocognitive deficits and greater risk for Alzheimer’s Disease (AD) being connected to declining testosterone levels among men (Zhang et al., 2024a) and falling estrogen levels among women (Kommaddi et al., 2023; Lopez-Lee et al., 2024).

To date, a categorical, holistic definition of sex has dominated both developmental and aging neuroscience investigations. Nonetheless, in recent years, this perspective has been challenged by findings suggesting that each human brain may be more accurately characterized as a tapestry of features that vary in frequency between sexes: some are most often detected in female brains, others, in male brains, while the remaining are equally likely to be observed in either sex (Joel et al., 2015). Shedding light on the medley of sex-(in)dependent characteristics unique to each brain and their trajectory of change across the life course could constitute a more effective strategy for personalizing detection and intervention paradigms for psychiatric and neurological conditions with sex-biased prevalence.

In late childhood and adolescence, relative alignment with female- vs male-typical functional connectivity (FC) patterns is linked to differential risk for mental health difficulties with sex-biased prevalence (i.e., mood vs aggressive behavior) (Petrican et al., 2025). Nonetheless, to our knowledge, it is yet to be determined whether relative similarity to female vs male-typical brain architecture is related to sex differences in cognitive-affective aging, specifically, severity of internalizing vs externalizing symptoms (Eme, 2016; Nathan et al., 2026; Yang et al., 2024) and vulnerability to neurodegenerative conditions, such as AD, which show distinct pathogenesis and progression among men vs women (Bachmann et al., 2023; Bourquard et al., 2023; Lopez-Lee et al., 2024; Ma et al., 2024). Linking the same regional patterns of alignment with sex-typical brain architecture to both aging-related neurodegeneration and internalizing-related problems could advance understanding of the well-documented link between depression and dementia in older adulthood (Dafsari and Jessen, 2020; Harerimana et al., 2022; Yang et al., 2023b Participants). Conversely, showing that neurodegeneration and internalizing problems relate to different patterns of sex-aligned FC would imply that distinct neurobiological processes underpin the differences between men and women observed in neurodegenerative vs internalizing conditions.

The importance of functional brain architecture in understanding the pathophysiology of neurodegenerative disorders, broadly, and of AD, in particular, is widely acknowledged. Consistent with the hypothesis that neurodegeneration spreads in an activity-dependent manner, regional FC strength has been shown to predict regional similarity in tau accumulation across the preclinical to dementia spectrum (Franzmeier et al., 2022; Franzmeier et al., 2019). Relatedly, the steeper decline observed in younger onset AD has been attributed to higher tau accumulation in frontoparietal regions, which accelerate brain-wide tau spreading through their extensive functional connections (Frontzkowski et al., 2022). FC patterns are not only a contributor, but also a robust correlate of progressive neurodegeneration, with association regions playing a critical role in the transition from normative to pathological aging. Specifically, in cognitively intact older adults, amyloid-beta deposition triggers excitation/inhibition imbalance in the default mode network (DMN), thereby amplifying excitability in medial temporal lobe (MTL) structures, a process which, in turn, initiates tau accumulation within the entorhinal cortex (Giorgio et al., 2023). More broadly, alterations along the canonical sensorimotor-association (S-A) axis, manifest as reduced dispersion, sensorimotor hypoconnectivity or association and limbic hyperconnectivity, track with cognitive deficits in mild cognitive impairment (MCI) and AD (He et al., 2023), as well as cumulative brain atrophy in dementia (including AD) (Brown et al., 2025).

Drawing on the reported relevance of FC to understanding sex-biased psychiatric vulnerabilities (Bangasser and Cuarenta, 2021; Dhamala et al., 2023; Petrican et al., 2025) and its well-documented contribution to AD onset (Brown et al., 2025; Franzmeier et al., 2022; Franzmeier et al., 2019; He et al., 2023), here, we leveraged a longitudinal, healthy aging dataset (aged 36 to 87 years) to test whether relative alignment with female- vs male-typical FC patterns correlates with future preclinical AD pathology and mental health problems, and whether any of the observed effects vary by sex and/or age (see Figure 1-A for an overview of the study procedure).

**Figure 1.**
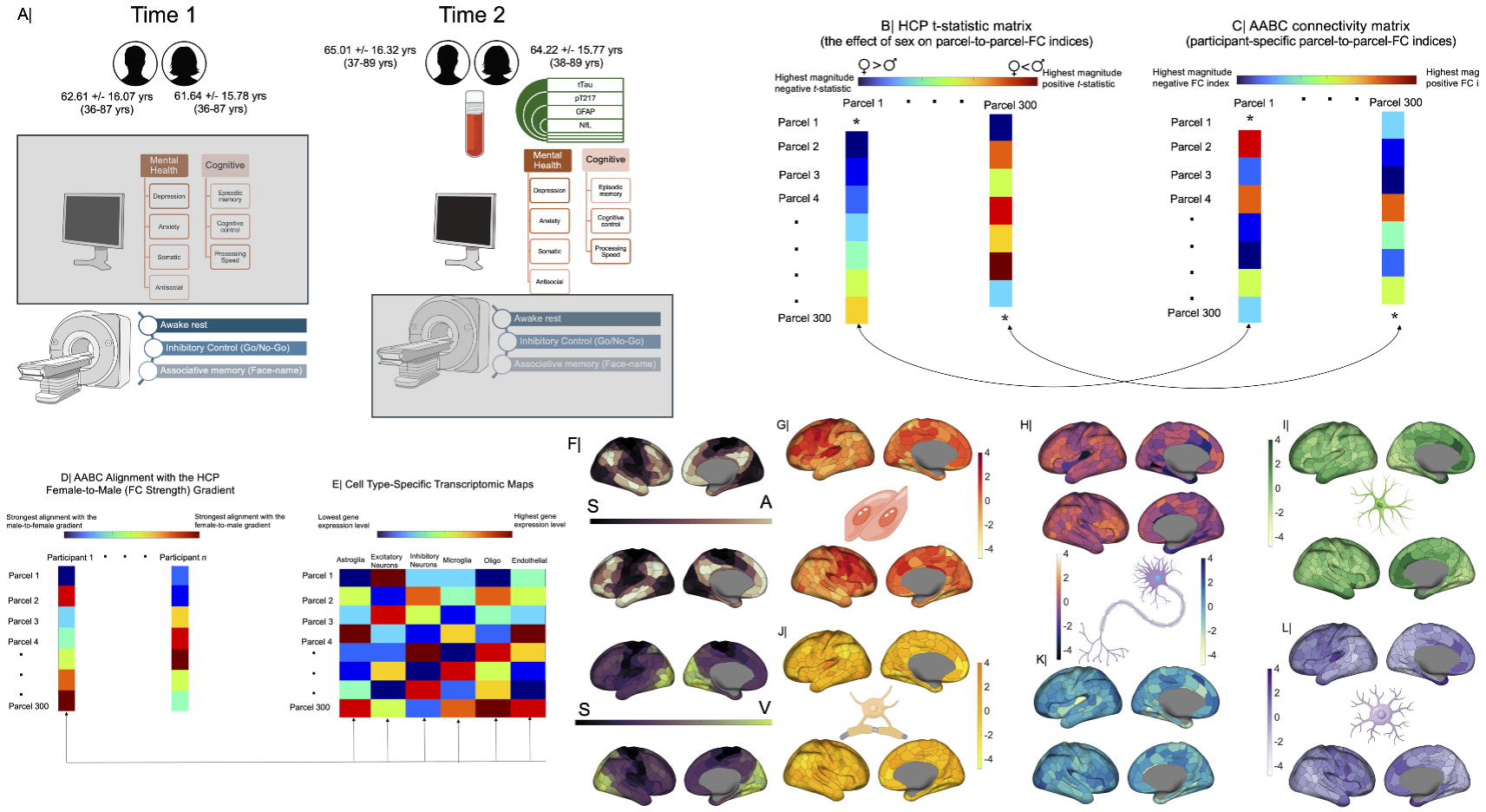
Overview of the study. Panel A: We analyzed two waves of mental health, cognitive, and neuroimaging data and one wave of brain injury/degeneration data from healthy aging adults in the AABC study (no brain injury/degeneration data was available at Time 1). Our objective was to investigate whether regional alignment with female- vs male-typical FC patterns during rest and/or task performance at Time 1 would relate to relate to mental health, neurodegeneration and cognitive performance at Time 2. All the corresponding multivariate models testing these relationships (i.e., CCAs 1 and 2) were adjusted for Time 1 mental health/cognition and Time 2 alignment with sex-typical FC (hence, the grey rectangles partially covering the corresponding variables in panel A). For details on covariate adjustment, see the Results corresponding to CCAs 1 and 2. For each AABC participant, relative alignment with female- vs male-typical FC patterns was computed as the rank-order correlation coefficient between each column of their own FC matrix (i.e., each parcel’s FC with each of the remaining parcels in the atlas) (panel C) and the corresponding column in the HCP *t*-statistic matrix (panel B). A positive correlation coefficient indicates that the regional FC strength profile from an AABC participant aligns with the female-to-male FC strength gradient from the HCP. A negative correlation coefficient indicates that the regional FC strength profile from an AABC participant aligns with the male-to-female FC strength gradient from the HCP. To estimate alignment with female- vs male-typical FC strength patterns along the two functional gradients (S-A, S-V) identified by Margulies et al. (2016) and six cell type-specific transcriptomic maps relevant to AD, we computed a rank order correlation between the matrix of sexual differentiation indices corresponding to the 300 parcels scrutinized (panel D) and each of the aforementioned eight spatial maps (panels E, F). Panel F: The canonical S-A (sensorimotor-to-association) and S-V (sensorimotor-to-visual) functional connectivity gradients. Panels G-L: Normalized bilateral expression of endothelial (panel G), excitatory neuronal (panel H), astroglial (panel I), oligodendrial (panel J), inhibitory neuronal (panel K) and microglial (panel L) genes based on the Schaefer atlas. Interpolation was used for parcels with non-robust gene expression values. Cognitive control = inhibitory control, cognitive flexibility, and working memory.

Relative alignment with female- vs male-typical FC was investigated along eight spatial brain maps relevant to understanding the spread of aging-related neurodegeneration. We thus examined the first two gradients (i.e., S-A and visual-sensorimotor [V-S]] described by Margulies et al. (Margulies et al., 2016), which capture sex and age differences in FC (Brown et al., 2025; He et al., 2023; Luo et al., 2024; Shanmugan et al., 2022) and the transcriptomic maps of genes regulating six cell types affected by AD, specifically, excitatory/inhibitory neurons, microglia, astroglia, oligodendrocytes and endothelial cells (Bourquard et al., 2023; Finze et al., 2023; Franzmeier et al., 2022; Lau et al., 2023; Richetin et al., 2020; Seidlitz et al., 2020). Our study pursued three interconnected objectives. First, we investigated whether, among aging women and men, relative alignment with female- vs male-typical FC patterns along the two canonical FC gradients and six cell type-specific transcriptomic maps outlined above would relate to subsequent accumulation of brain injury/degeneration markers relevant to AD and greater severity of mood symptoms. Alignment with sex-typical FC was examined during awake rest and during performance of two tasks (i.e., inhibitory control, associative memory) which are sensitive to aging-related decline and internalizing-related problems (Kamp et al., 2026; Keller et al., 2019; Li et al., 2005; Spreng and Turner, 2019). Second, we investigated whether the sex-aligned FC patterns identified at point (1) would show longitudinal associations with processing speed, episodic memory and/or cognitive control, mental processes that tend to be affected by aging and internalizing symptomatology (Colwell et al., 2022; Damgaard et al., 2025; James et al., 2021; Spreng and Turner, 2019). Third, gonadal hormones drive neural sexual differentiation (Gegenhuber et al., 2022; Küchenhoff et al., 2024) and, thus, likely sex differences in brain architecture. Therefore, we examined the overlap between the six cell type-specific transcriptomic maps, interrogated for relative alignment with female- vs male-typical FC patterns (see point 1), and the transcriptomic maps of genes regulating sex steroid synthesis and their receptor functions. A robust overlap would raise the possibility that neurodegenerative and/or internalizing risk could be related to regional sex hormone modulation of cell type-specific activity (Lee et al., 2026; Nathan et al., 2026).

## Methods and Materials

The main methodological features of the study are listed below and detailed in the Supplemental Information (SI).

### Participants

We analysed a publicly available community dataset (see Figure 2 for the selection criteria). All participants were healthy aging adults from the Aging Adult Vulnerability and Resiliency in the Aging Adult Brain Connectome (AABC, (Bookheimer et al., 2019)) project (N = 353, M_age_ = 62.04 years, SD_age_ = 15.77 years [range 36 to 87 years] at wave 1, 208 women) who provided data at two waves across an average of 2.5 years (SD = 1.28 years, range 1 to 7 years). There was only one testing session at each wave. Independent-samples Mann-Whitney U tests confirmed that there were no sex differences in age at either wave (both *p*s > .59). Most participants were predominantly right-handed and self-identified as White. The delay between the two visits was included as a covariate in all the analyses reported under “Results”.

**Figure 2.**
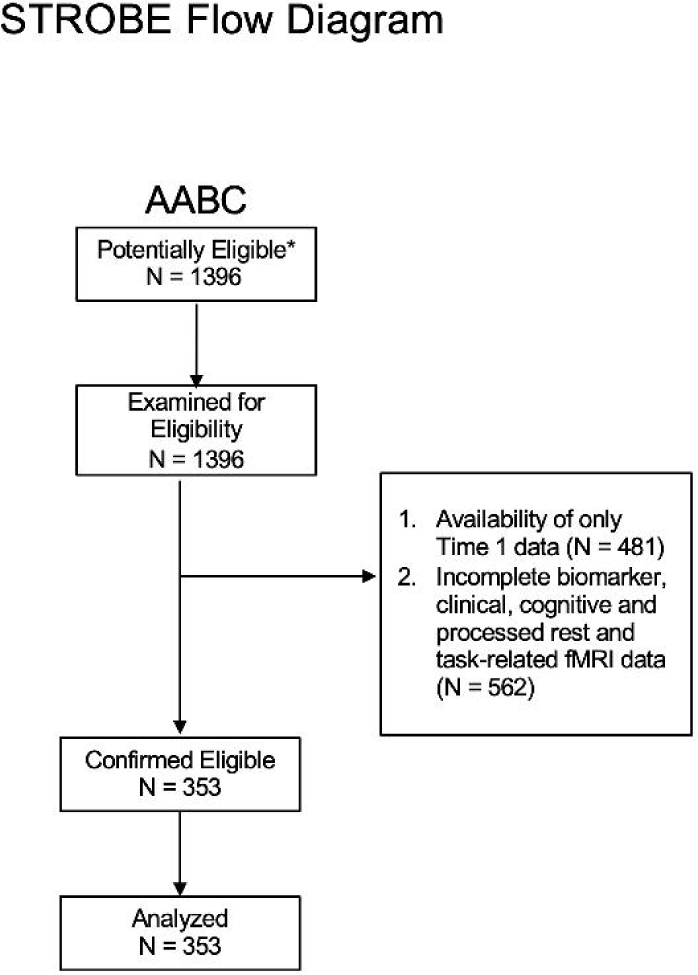
STROBE flowchart describing sample selection for the AABC sample.

### Mental Health Assessments (Time 1 and Time 2)

Internalizing (depression, anxiety, somatic complaints) and externalizing (antisocial personality) symptoms were assessed via the corresponding DSM-oriented scales from the Adult Self-Report Scale (ASR, (Achenbach, 2009) (for adults under 60 years) or the Older Adult Self-Report Scale (OASR, (Achenbach et al., 2004) (for adults aged 60 years and above). Although our research questions focused on internalizing symptoms, we also assessed externalizing problems in order to determine the specificity of any internalizing-FC associations.

### Cognitive Functioning (Time 1 and Time 2)

We included five fluid cognitive tasks on which performance tends to show a robust decline in both healthy and pathological aging, as well as in internalizing conditions across the life course (Colwell et al., 2022; Damgaard et al., 2025; James et al., 2021; Spreng and Turner, 2019). Specifically, we analyzed scores on episodic memory (Picture Sequence Task), inhibitory control (Flanker Task), working memory (List Sorting Task), processing speed (Pattern Comparison Task) and cognitive flexibility (Dimensional Card Sorting task). Detailed descriptions of these tasks have already been published (Bookheimer et al., 2019) (for a summary, see SI). All the reported analyses controlled for education level as a proxy for crystallized cognitive capacity and socioeconomic status.

### Brain Injury Markers (Time 2 only)

We analyzed the blood concentrations (pg/mL) of four markers of neurodegeneration and neural injury, neurofilament light chain (NfL, a marker of axonal injury, M = 129.26, SD = 108.42, range: 30.74-1461.52), glial fibrillary acidic protein (GFAP, a marker of astrocytic activation, M = 59.47, SD = 36.46, range: 11.57-231.85), total Tau (tTau, M = 28.66, SD = 14.05, range: 11.19-153.04) and Phospho-Tau T217 (pT217, M = 3.88, SD = 3.26, range: .91-31.21). These biomarkers, which are part of the AABC Data Release 2.0, have been quantified based on blood samples collected early morning after 8 hours of fasting (see Supplementary Figure 2 in (Bookheimer et al., 2019)). They had been implicated in normative aging (Babulal, 2025; Bookheimer et al., 2019) and in the accelerated neurobiological aging triggered by environmental stressors (e.g., viral agents,(Wood et al., 2025)).

### Gene Expression Profiles

Brain-wide expression of cell type-specific (i.e., astroglial, neuronal [excitatory, inhibitory], microglial, oligodendrial, endothelial) (Seidlitz et al., 2020) and sex hormone (i.e., androgen, estrogen, progesterone receptor family, sex steroid synthesis; see Table S1) genes (Küchenhoff et al., 2024) was estimated in abagen (Markello et al., 2021) from the microarray data of the six donors released by the Allen Brain Institute. Because gene expression is symmetrical across hemispheres, our analyses were based on transcriptomic data mirrored across hemispheres and used interpolation for missing data. However, in supplemental tests, we verified that all the results hold if we used only left hemisphere ROIs—to which all six donors contributed--with no interpolation for missing data (see Supplemental Results [SI]).

### fMRI Data Acquisition and Processing (Time 1 and Time 2)

The resting state and task-related (face-name associative memory, inhibitory control, (Bookheimer et al., 2019)) functional neuroimaging data used in the present report had been cleaned with the Human Connectome Project (HCP) Preprocessing Pipelines (Glasser et al., 2013). This is the same preprocessing pipeline applied to the HCP young adult group who served as a reference for the sex-typical FC gradient analyses described below. The acquisition and preprocessing protocols for the AABC and HCP samples had been already published (Harms et al., 2018; Smith et al., 2013) and we summarize them in the SI.

### Sex-Typical FC Gradient

Female- vs male-typical FC patterns were estimated based on the resting state functional neuroimaging data from healthy young adult HCP participants (N = 336, 159 women, aged 22-30 years) for whom sex differences in brain function, stemming from the interplay of sex chromosomal and gonadal hormonal effects (Ahmed et al., 2008; Gilmore et al., 2018; Hines et al., 2015; Pfeifer and Allen, 2021; Piekarski et al., 2023; Sato et al., 2004), were expected to have peaked (Petrican et al., 2025). Use of the AABC participants as a reference for defining the female-to-male FC gradient would have been suboptimal for two key reasons. One is the steeper physiological aging slope observed among women (Lopez-Lee et al., 2024), which means that the definition of the female-to-male gradient could have confounded aging-related and sex-related differences. The second reason for opting against the AABC as a reference group is the aging-related decline in gonadal hormones (Nathan et al., 2026), which would have likely impacted the magnitude of any sex differences in FC.

The main analyses were conducted on a widely used 300-parcel functional brain atlas (Schaefer et al., 2018) and replicated with a second, independently defined atlas (see SI). To identify sex-typical connectivity patterns in the HCP, we regressed each of the 44850 parcel-to-parcel FC indices against sex and a number of confounds which had been linked to individual differences in FC and on which complete matching between sexes is not feasible, specifically, age, income, employment status, race, handedness, average in-scanner motion, intelligence and psychopathology. For each parcel-to-parcel FC index, the *t*-statistic corresponding to “sex” was taken as a measure of the robustness with which the respective FC index differentiates between men and women. We opted to focus on the t-statistic, rather than the beta weight corresponding to “sex”, in order to safeguard against prioritising FC indices that show large sex differences in a subsample, but not the entire HCP group (i.e., FC indices with high absolute value beta weights, but also high standard errors, suggestive of greater between-individual variability and, thus, lower robustness in distinguishing between sexes). Nonetheless, we verified that the *t*-statistic and corresponding beta weight value associated with “sex” for a given parcel-to-parcel FC index were very strongly correlated (median *r* of .96, range: .79-.995), as expected given the intentional selection of a relatively homogenous group of HCP participants to act as reference for the sex-related FC gradient analyses. We further examined whether the sex-related FC effects observed in the HCP sample (i.e., the *t*-statistic values corresponding to “sex”) had been biased by the skewness of the covariates entered in the original regression analyses. To this end, we ran a Spearman’s partial correlation analysis between sex and each of 44850 FC estimates, while adjusting for all the nuisance variables entered in the corresponding regression analysis. In line with the interpretation that data skewness was unlikely to bias the sex-relevant regression results, the correlation between the sex-related *t*-statistic from the original regression analysis and the sex-related Spearman’s rho for each of the 44850 FC estimates was very high (median *r* of .97, range: .85-.99).

### Alignment with Sex-Typical FC Patterns (Time 1 and Time 2)

For each AABC participant, relative alignment with female- vs male-typical FC patterns was computed as the rank-order correlation coefficient between each column of their own FC matrix (i.e., each parcel’s FC with each of the remaining parcels in the atlas) and the corresponding column in the HCP *t*-statistic matrix. Because the HCP *t*-statistic matrix is based only on young adult FC profiles, anchored at one end in female prototypicality and, at the other, in male prototypicality, both positive negative rank order correlation coefficients indicate similarity to a young adult brain profile. Thus, for each individual, parcels that align more strongly with the HCP male-typical (rather than the female-typical) FC strength pattern have a higher magnitude positive coefficient. Complementarily, parcels that align more strongly with the HCP female-typical (rather than the male-typical) FC strength pattern have a higher magnitude negative coefficient. Figure 1 contains a representation of the sex-related FC gradient analyses in the HCP sample and of the sex-related FC alignment tests in the AABC sample. We replicated all the results with HCP sexual differentiation models further adjusted for brain size and individual differences in female hormonal state (see Supplemental Results [SI]).

### Validation of the HCP Sex-Typical FC Gradient in the AABC Sample (Time 1 and Time 2)

Both HCP sexual differentiation models were validated in the AABC sample by confirming that the male participants in both samples showed relatively stronger alignment in FC with the HCP men (rather than the HCP women), whereas the female participants showed stronger alignment in FC with the HCP women (rather than the HCP men). Furthermore, across both samples (AABC, HCP), association (relative to unimodal) regions showed stronger resting state (i.e., intrinsic) alignment with male FC patterns among men and stronger resting state (i.e., intrinsic) alignment with female FC patterns among women (see Supplemental Methods). This pattern of within-individual variability in alignment with sex-typical FC dovetails with previous reports that association regions are most effective at classifying individuals based on sex (Shanmugan et al., 2022). Taken together, these analyses attest that FC patterns that differentiate between the young HCP women and men can similarly differentiate between the older AABC women and men.

### Statistical Analysis

We relied primarily on canonical correlation analysis (CCA), a multivariate data-driven technique which explains commonalities in two sets of variables by creating linear combinations, dubbed “variates”, from the observed variables in each set (Mihalik et al., 2022). Our analytic strategy is grounded in evidence that both normative and pathological lifespan development emerge from the interactions of multiple functional systems (Gao, 2025; Graham et al., 2021; Insel et al., 2010; Williams et al., 2024; Williams et al., 2016). All the CCA models underwent 10-fold cross-validation (i.e., the model discovery and testing were performed on non-overlapping subsets of the same sample). Permutation testing (100,000 samples) helped determine statistical significance, whereas bootstrapping (100,000 samples) established the reliability of the model estimates (95% confidence intervals [CIs]). We conducted three multivariate analyses (i.e., CCAs) for which we ran the technical replications reported in the SI. Statistical significance in the discovery and corresponding cross-validation (aka “simulated replication”) CCAs was assessed only for the variate pair explaining the highest amount of covariance between the two sets of observed variables in each of the three discovery analyses. There was no need to correct across the three CCAs because each analysis considered a different type of data.

## Results

Below we label as “statistically significant” or “statistically reliable” only the results that meet these criteria across all the control tests (“technical replications”) reported in the SI.

### CCA 1: Alignment with Sex-Typical Functional Brain Profiles Relates to Future Depression and Neurodegeneration

We started by examining whether individual differences in neurodegeneration markers and psychiatric symptoms are linked to earlier alignment with sex-typical FC patterns along the two functional gradients (S-A, S-V) identified by Margulies et al. (2016) and six cell type-specific transcriptomic maps relevant to AD ([excitatory/inhibitory] neurons, astroglia, microglia, oligodendria, endothelial cells). To this end, for each AABC participant, we computed a rank order correlation between the matrix of sexual differentiation indices corresponding to the 300 parcels scrutinized (see Methods above) and each of these eight spatial maps. Positive coefficients indicated greater alignment with male-typical (rather than female-typical) FC patterns in transmodal (S-A gradient), visual (S-V gradient) or in brain regions enriched for a cell type specific gene set. Negative coefficients indicated that transmodal (S-A axis), visual (S-V axis) or areas enriched for a cell type specific gene set show relatively greater similarity to female-typical (rather than male-typical) FC patterns. To probe mental context specificity, the eight alignment indices were estimated during both awake rest and task (i.e., inhibitory control, associative memory) performance.

Next, we linked via CCA Time 2 chronological age, psychopathology (i.e., depression, anxiety, somatic complaints, antisocial personality) and neurodegeneration (i.e. tTau, pT217, GFAP, Nfl) to sex and Time 1 alignment with sex-typical FC along the eight spatial maps listed above. Age and sex were placed in the same set as the variables with which they are most strongly connected (i.e., neurodegeneration vs neural sexual differentiation) in order to avoid biasing the extraction of the CCA variate pair (e.g., avoid extracting a CCA pair that only reflects age-related neurodegeneration). Because we controlled for the delay between the two visits (see below), the interpretation of the age-related results is identical if age is included in the Time 1 or Time 2 variable set.

This analysis yielded one CCA mode which was successfully cross-validated (*r*_cv_ = .21, permutation-based *p* = 15 × 10^-5^, Figure 3-E) after adjusting for the corresponding Time 2 alignment with sex-typical FC and Time 1 psychopathology indices, in-scanner motion, scanner type, delay between the two testing sessions, education, race, handedness and number of medications taken at each of the two time points. The cross-validated variate pair distinguished between older men showing greater neurodegeneration and younger women experiencing more severe depression at Time 2. These two sex-dependent profiles were associated with region- and context-dependent patterns of alignment with sex-typical FC at Time 1. Specifically, younger women who were more severely depressed at Time 2 showed stronger Time 1 alignment with female-typical (rather than male-typical) connectivity patterns in regions enriched for microglial (rest, cognitive control performance), endothelial (rest, associative memory performance) and oligodendrial (cognitive control performance) genes (Figure 3-A, C). Complementarily, older men with higher GFAP and pT217 blood concentration at Time 2 demonstrated greater alignment with male-typical (rather than female-typical) connectivity patterns in the same brain areas at Time 1 (Figure 3-B, D). From an architectural standpoint, the resting state sexual differentiation patterns were expressed more strongly in unimodal, rather than transmodal, specifically, sensorimotor rather than visual areas. In a similar vein, during cognitive control performance, patterns of sexual differentiation related to later neurodegeneration or depression were more salient in sensorimotor rather than visual areas.

**Figure 3.**
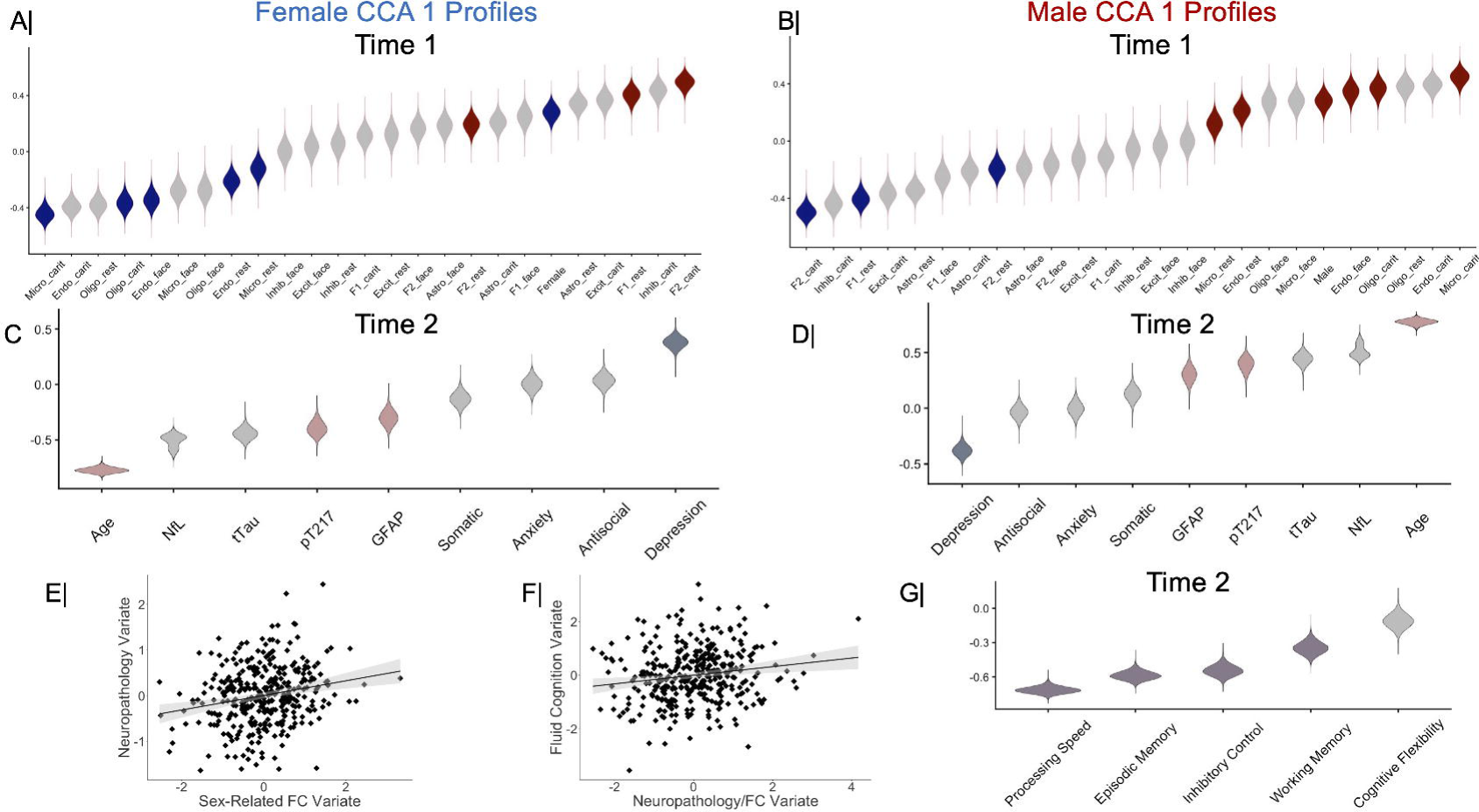
Cross validated variates from CCAs 1 and 2 describing the inter-relationships among context- and sex-dependent functional connectivity patterns, neuropathology and cognitive performance. The violin plots in panels A-D and G depict the distribution of correlation coefficients across the 100,000 bootstrap samples from the cross-validation procedure for CCA 1 (A-D) and CCA 2 (G). In these panels, the y-axes represent the cross-validated canonical loadings for each CCA. Gray colored violin plots indicate non-robust canonical loadings (based on the bootstrapped 95% CIs from the cross-validation procedure across all the control conditions scrutinized [atlas x laterality x HCP female-to-male FC gradient]). In panels A and B, blue violin plots indicate female (rather than male)-aligned FC patterns along the S-A (F1) gradient, S-V (F2) gradient or in brain regions enriched for cell-type specific genes. Complementarily, red violin plots indicate male (rather than female)-aligned FC patterns along the same spatial maps. The CCA 1 female variates pictured in panels A and C were generated by multiplying the corresponding CCA 1 male variates by (−1). In other words, the female and male CCA 1 variates represent the two sides of the same variate pair pictured separately only to facilitate interpretation. Panels E and F contain the scatter plot describing the linear relationship between the cross-validated variates from CCA 1 (E) or CCA 2 (F). In these panels, the gray shaded area represents the 95% CI of the corresponding correlation coefficient. CI = confidence interval. FC = functional connectivity. F1 = sensorimotor-to-association gradient. F2 = sensorimotor-to-visual gradient. Astro = astroglia. Excit = excitatory neurons. Inhib = inhibitory neurons. Micro = microglia. Oligo = oligodendrocytes and oligodendrocyte precursor cells. Endo = endothelial cells. Carit = inhibitory control task. Face = face-name association task.

**Figure 4.**
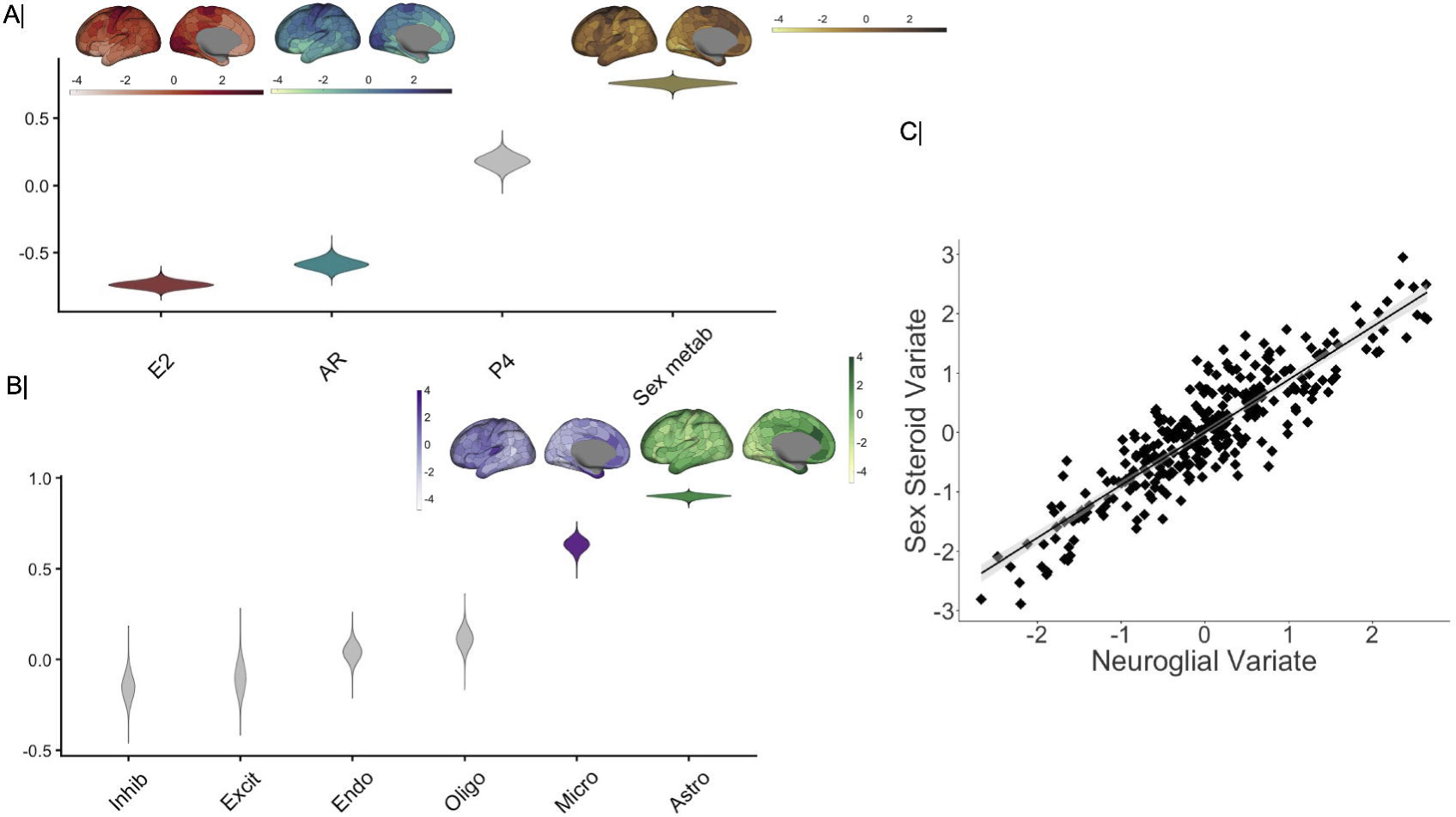
Cross validated variates from CCA 3 describing the relationship between cell type-specific and sex steroid-related gene expression. The violin plots in panels A and B depict the distribution of correlation coefficients across the 100,000 bootstrap samples from the cross-validation procedure. In these panels, the y-axes represent the cross-validated canonical loadings. Gray colored violin plots indicate non-robust canonical loadings (based on the bootstrapped 95% CIs from the cross-validation procedure across all the control conditions scrutinized [atlas x laterality x HCP female-to-male FC gradient]). Panel C contains the scatter plot describing the linear relationship between the cross-validated variates from CCA 3. In this panel, the gray shaded area represents the 95% CI of the corresponding correlation coefficient. CI = confidence interval. FC = functional connectivity. F1 = sensorimotor-to-association gradient. F2 = sensorimotor-to-visual gradient. AR = androgen receptor. E2 = estradiol (refers here to estrogen receptor genes). P4 = progesterone (refers here to progesterone receptor family genes). Metab = metabolism. Astro = astroglia. Excit = excitatory neurons. Inhib = inhibitory neurons. Micro = microglia. Oligo = oligodendrocytes and oligodendrocyte precursor cells. Endo = endothelial cells.

### CCA 2: Neurodegeneration/Depression-Relevant Alignment with Sex-Typical Functional Brain Profiles Relates to Cognitive Performance

Next, we sought to confirm the behavioural relevance of the Time 1 sex-dependent FC and Time 2 neurodegeneration/psychopathology variates described above. To this end, we linked them via CCA to Time 2 (i.e., neurodegeneration-concurrent) scores on mental functions which tend to decline with aging and internalizing symptomatology, specifically, inhibitory control, cognitive flexibility, working memory, processing speed and episodic memory. The emergent CCA mode was successfully cross-validated (*r*_cv_ = .16, permutation-based *p* = .005, 95% CI= [.04; .28], Figure 3-F) after adjusting for all the confounds entered in the cross-validation of the CCA 1 results (see prior section) and the corresponding Time 1 cognitive performance scores. The results indicated that poorer inhibitory control, episodic memory, processing speed and working memory at Time 2 are robustly related to the CCA 1 variates, specifically, the Time 2 higher neurodegeneration/lower depression variate (loading_CV_ = .99, 95% CI= [.99; 1.0]) and the associated male FC variate at Time 1 (loading_CV_ = .31, 95% CI= [.20; .40], Figure 3-B, D, G).

### CCA 3: Robust Overlap in the Transcriptomic Maps of Neuroinflammation- and Sex Steroid-Relevant Genes

The results of CCA 1 indicated that areas enriched for microglial, endothelial and oligodendrial genes show sex-related FC variations which may relate to vulnerability to neurodegeneration vs internalizing symptoms (i.e., depression). To further explore this possibility, we drew on evidence that the most sexually differentiated brain regions tend to be enriched for sex steroid genes (Küchenhoff et al., 2024). Consequently, in CCA 3, we examined the spatial overlap between the transcriptomic maps of the cell type-specific genes from CCA 1 and the transcriptomic maps of the four sex hormone gene types identified by (Küchenhoff et al., 2024).

We included all six cell type gene sets because we sought to determine the specificity of any observed associations with the sex steroid genes. The cross-validated mode emerging from CCA 3 (*r*_cv_ = .89, *p*_spin_ = 10^-5^, Figure 5-C) reflected the intersection between the transcriptomic maps of astroglial and microglial genes and of genes regulating sex steroid metabolism (Figure 5-A, B). The results of this analysis remained unchanged after eliminating the 8 sex hormone genes with preferential expression in specific cell types. Taken together, the results of CCAs 1 and 3 raise the possibility that sex hormone-related modulation of microglial activity may be relevant to understanding the link between sex-dependent FC patterns and later neurodegeneration among men or depression among women.

## Discussion

Extending prior literature on the role of the brain’s functional architecture in internalizing symptoms, as well as AD onset and progression (Bangasser and Cuarenta, 2021; Dhamala et al., 2023; Franzmeier et al., 2019; Frontzkowski et al., 2022; Petrican et al., 2025), we provide novel evidence that regional alignment with female- vs male-typical FC patterns constitutes a sex- and age-dependent longitudinal correlate of brain injury/degeneration vs depression. Specifically, among older men, deviance from male-typical brain architecture, manifest as stronger resting state alignment with male-typical FC patterns in unimodal, rather than association, areas (Shanmugan et al., 2022) correlated with higher concentrations of pT217 and GFAP, two key biomarkers of longitudinal neural atrophy and fluid cognitive decline in aging and (pre-)clinical AD (Ferrari-Souza et al., 2022; He et al., 2025; Heneka et al., 2025; Huang et al., 2026; Zhang et al., 2024b), which track with genetic risk for AD from midlife onwards (i.e., GFAP, (Freudenberg-Hua et al., 2025)). Deviance from female-typical FC along the same S-A axis correlated with (subclinical) depression severity among younger women. Both the male and female FC profiles spanned regions enriched for microglial, endothelial or oligodendrial genes, with some evidence of context-specificity (i.e., sex FC-microglial maps: rest and cognitive control performance; sex FC-endothelial maps: rest and associative memory performance; sex FC-oligodendrial maps: cognitive control performance). Broadly, these results resonate with prior reports that glial cells undergo a host of sex-dependent changes during brain aging (Zhang et al., 2024c DNA damage). These changes are conducive to sex-dependent functionalities, which could potentially explain their differential involvement in female depression vs male AD-linked brain pathology.

The FC-transcriptomic map associations related to later neurodegeneration among older men fit nicely with the well-documented role of microglia, oligodendria and endothelial cells in normative aging and the transition to AD pathophysiology (Bourquard et al., 2023; d’Errico et al., 2022; Kosoy et al., 2025; Lau et al., 2023; Reid et al., 2025; Zhang et al., 2024c DNA damage). There is substantial evidence that accumulation of senescent glial cells drives neuroinflammation, a typical feature of normative aging which can nonetheless heighten risk for AD by enhancing systemic inflammation and accelerating brain aging (Basurco et al., 2024; Zhang et al., 2024c DNA damage). Among glial cells, microglia are regarded as the main drivers of central inflammation in both normative late life development and pathological conditions (Groh et al., 2025; Singh et al., 2023). Specifically, microglia are thought to facilitate spread of beta-amyloid pathology to previously unaffected brain regions (d’Errico et al., 2022), while accumulation of senescent microglia allegedly forecasts the transition from healthy cognition to MCI and later AD (Lau et al., 2023).

In our analyses, brain regions enriched for microglial genes showed also stronger expression of genes regulating sex steroid metabolism. Sex steroids have been implicated in neural sexual differentiation (Gegenhuber et al., 2022), including expression of sexually dimorphic phenotypes in adulthood (Sato et al., 2004). Therefore, it is possible that greater androgen availability in areas enriched for microglial genes may be one of the mechanisms driving the stronger alignment with male-typical (rather than female-typical) FC observed among older men at risk for later neurodegeneration. Recent reports suggest that androgens can suppress an inflammatory response in the microglia (Lee et al., 2026). Therefore, in our study, male-biased FC patterns in regions enriched for microglial genes could reflect an initial compensatory response through which potential increases in regional androgen availability could dampen early central inflammatory responses among currently healthy older men who are on-course for developing later neurodegeneration. Our proposed compensatory account is also compatible with the fact that greater alignment with male-typical (rather than female-typical) FC in microglia-rich areas was linked to a relatively constrained profile of neurodegeneration and functional deficits (i.e., impaired processing speed, episodic memory, inhibitory control, working memory, but no robust deficits in cognitive flexibility and reduced depression) among older men who did not receive a diagnosis of either MCI or AD over the scrutinized period.

A conceptually similar interpretation could apply to the complementary longitudinal association between stronger alignment with female-typical (rather than male-typical) FC in microglia-rich regions and more severe depression, as observed among the younger AABC women. Indeed, declining estradiol (E2) levels during the menopausal transition have been robustly linked to increased vulnerability to depression and greater neuroinflammation, the latter due to the anti-inflammatory effects of E2 on microglial activation (Nathan et al., 2026). Given the contribution of gonadal hormones to sexual differentiation (Gegenhuber et al., 2022; Küchenhoff et al., 2024), the female-biased FC patterns observed in microglia-rich regions could stem from a compensatory increase in E2 regional availability which is triggered to attenuate central inflammatory responses among women on course for more severe depression. In the AABC sample, this reasoning is compatible with the finding that younger women who showed greater baseline alignment with female-typical (rather than male-typical) FC in microglia-rich areas demonstrated superior cognitive performance at the follow-up despite the greater severity of their depressive symptoms (which nonetheless remained at subclinical levels).

There is suggestive evidence that early life adversity can set in motion a suboptimal lifespan developmental sequence, which is punctuated by both externalizing and internalizing psychopathology (McLaughlin et al., 2020) and may culminate in dementia in later adulthood (Radford et al., 2017; Tani et al., 2020). The detrimental impact of early life adversity on aging outcomes is augmented through ongoing exposure to stressors {Lou, 2025, childhood adversity; Zhou, 2025, stressful life events). Paradoxically, though, in some instances, negative childhood events can inoculate against later experienced adversity and foster psychological resilience in middle to older adulthood (Zong et al., 2025). Understanding the sex-differential neurobiological mechanisms though which early life adversity may heighten or protect against vulnerability to psychopathology and dementia in older adulthood, as well as the modulating influence of cognitive reserve (Cabeza et al., 2018 reserve and compensation) and social support, would substantially advance efforts to personalize interventions for lifelong mental health.

Our investigation has several limitations which could nonetheless pave the way for new research directions in the future. First, our analyses were based on two waves of data from a predominantly White sample. More extensive assessments spanning several timescales (e.g., on the order of days vs decades) would provide further insights into the effects herein reported. Second, neither genotyping nor epigenetic data were available. Future studies incorporating cell-type specific (e.g., microglia vs astrocytes, (Yang et al., 2023a)) genetic risk scores for AD and MDD in addition to epigenetic (e.g., DNA methylation-based) and physiological aging measures could help identify more precise intervention targets for both dementia and mood-related disorders. Third, cross-species experimental manipulations of regional sex steroid availability and neuroinflammatory responses in specific cell types (i.e., microglia, astroglia) would help shed further light on the potential modulatory effect of gonadal hormones on neuron-glia interactions (Bourquard et al., 2023; Gu et al., 2025; Tournier et al., 2024; Vila-Castelar et al., 2025; Zhang et al., 2024c DNA damage).

In sum, we found evidence that, in middle to older adulthood, internalizing symptoms and neurodegeneration show longitudinal correlations with complementary, rather than identical, patterns of deviance from sex-typical brain architecture, which are differentially expressed among men and women as a function of age. Both profiles overlapped areas enriched in microglial genes and genes regulating sex steroid metabolism, in line with evidence that the interplay between gonadal hormones and neuro-inflammation is key to understanding aging-related differences in cognition and affect. Taken together, our findings attest to the value of integrating multilevel assessments of brain architecture, physiology and gene expression in order to fine-tune precision medicine approaches across the life course.

## Data statement

The raw data are available https://balsa.wustl.edu (AABC) and at https://db.humanconnectome.org (HCP) upon completion of the relevant data use agreements. The AABC data used in this report came from Data Release 2.0.

## Code availability

We used already existing code, as specified in the main text with links for free download.

## Conflict of interest

The authors declare no competing interests.

## Acknowledgments

Data used in the preparation of this article were obtained from the Aging Adult Vulnerability and Resiliency in the Aging Adult Brain Connectome (AABC) project, the Human Connectome Project in Aging (HCP-A), and the Human Connectome Project, WU-Minn Consortium (Principal Investigators: David Van Essen and Kamil Ugurbil; 1U54MH091657; funders: the 16 NIH Institutes and Centers that support the NIH Blueprint for Neuroscience Research and the McDonnell Center for Systems Neuroscience at Washington University). The AABC (U19AG073585) and HCP-A (U01AG052564) were funded by the National Institute of Aging of the National Institutes of Health. HCP-A was further supported by funds provided by the McDonnell Center for Neuroscience at Washington University in St. Louis. This manuscript reflects the views of the authors and may not reflect the opinions or views of the NIH, AABC, HCP-A, or HCP consortium investigators.

## Notes

### Competing Interest Statement

The authors have declared no competing interest.

